# Activation of SEDS-PBP cell wall synthases by an essential regulator of bacterial division

**DOI:** 10.1101/453415

**Authors:** Patrick J. Lariviere, Christopher R. Mahone, Gustavo Santiago-Collazo, Matthew Howell, Allison K. Daitch, Rilee Zeinert, Peter Chien, Pamela J. B. Brown, Erin D. Goley

**Affiliations:** Department of Biological Chemistry, Johns Hopkins University School of Medicine, Baltimore, MD 21205, USA.; Division of Biological Sciences, University of Missouri, Columbia, MO 65211, USA.; Department of Biochemistry and Molecular Biology, University of Massachusetts Amherst, Amherst, MA 01003, USA.

## Abstract

Bacterial growth and division require insertion of new peptidoglycan (PG) into the existing cell wall by PG synthase enzymes. Emerging evidence suggests that many PG synthases require activation to function, however it is unclear how activation of division-specific PG synthases occurs. The FtsZ cytoskeleton has been implicated as a regulator of PG synthesis during division, but the mechanisms through which it acts are unknown. Here we show that FzlA, an essential regulator of constriction in *Caulobacter crescentus*, links FtsZ to PG synthesis to promote division. We find that hyperactive mutants of the PG synthases FtsW and FtsI specifically render *fzlA*, but not other division genes, non-essential. However, FzlA is still required to maintain proper constriction rate and efficiency in a hyperactive PG synthase background. Intriguingly, loss of *fzlA* in the presence of hyperactivated FtsWI causes cells to rotate about the division plane during constriction and sensitizes cells to cell wall-specific antibiotics. We demonstrate that FzlA-dependent signaling to division-specific PG synthesis is conserved in another α-proteobacterium, *Agrobacterium tumefaciens.* These data establish that FzlA links FtsZ to cell wall remodeling, serving both to activate and spatially orient PG synthesis during division. Overall, our findings support the paradigm that activation of SEDS-PBP PG synthases is a broadly conserved requirement for bacterial morphogenesis.

---

Bacterial division is driven by the insertion of new cell wall material at midcell in a tightly regulated manner, allowing for determination of cell shape and maintenance of envelope integrity^1,2^. The cell wall is made of peptidoglycan (PG), a meshwork consisting of glycan strands crosslinked by peptide stems^3,4^. PG synthesis requires the coordination of glycan polymerization and peptide crosslinking by either coupled monofunctional glycosyltransferases (GTases) and transpeptidases (TPases), or bifunctional enzymes that contain both activities, with these proteins being more generally referred to as PG synthases^1^.

Monofunctional PG synthase pairs have been implicated as the primary synthetic enzymes of the elongation (elongasome) and division (divisome) machineries. A paradigm has been proposed whereby a shape, elongation, division, and sporulation (SEDS) family GTase is functionally coupled to a penicillin binding protein (PBP) TPase, which together facilitate cell wall synthesis^5–7^. Through characterization of the elongation-specific PG synthases RodA and PBP2 in *Escherichia coli*, it has been postulated that SEDS-PBP enzymes require activation to function^5^. Specifically, mutations in RodA or PBP2 that increase GTase activity *in vitro* and PG synthesis in cells render other components of the elongasome non-essential, arguing that their normal function is to activate the RodA-PBP2 complex^5^. Intriguingly, analogous mutations in the division-specific SEDS-PBP enzymes, FtsW and FtsI, allow cells to constrict faster than normal^8^, suggesting that these mutations promote formation of an activated PG synthase complex^5,9^. However, it is unclear precisely how SEDS-PBP activation normally occurs during division.

Recent studies have established that the conserved cytoskeletal protein FtsZ^10,11^, which recruits the division machinery to a ring-like structure at midcell^12–14^, is coupled to PG synthesis activation during division. In multiple organisms, the C-terminal linker domain of FtsZ was found to be required for regulating cell wall integrity^15–17^ and shape, as well as PG chemistry^16,18^. Moreover, in *E. coli* and *Bacillus subtilis*, FtsZ dynamics were demonstrated to drive PG synthase dynamics in both organisms, as well as division site shape in *E. coli* and constriction rate in *B. subtilis*^19,20^. Collectively these data indicate that, at least in some organisms, FtsZ acts as a “dynamic scaffold” or “dynamic activator” of PG synthesis likely impinging on FtsWI. However, the signaling pathway connecting these two endpoints remains unresolved.

We previously demonstrated that an essential FtsZ-binding protein, FzlA^21^, is required for division and regulates the rate of constriction in the α-proteobacterium *Caulobacter crescentus*^22^. Mutations in FzlA with diminished affinity for FtsZ were found to have slower constriction rates and altered cell pole shape, indicative of reduced PG synthetic activity during division^22^. We therefore postulated that FzlA facilitates a link between FtsZ and PG synthesis by serving as an upstream activator of PG synthases and, here, set out to test this hypothesis.

## Results

### *fzlA* lies upstream of *ftsWI* in a PG synthesis pathway

We reasoned that if FzlA impacts constriction through PG synthases, it likely acts on the division-specific SEDS family GTase FtsW and/or the monofunctional PBP TPase FtsI. To assess if FzlA activates FtsWI, we leveraged fast-constricting strains containing hyperactive mutant variants of FtsI and/or FtsW termed *ftsW**I\**^8,9^ and *ftsW\**^9^*. ftsW**I\** bears the mutations F145L and A246T in FtsW and I45V in FtsI, whereas *ftsW\** contains only the FtsW A246T mutation^9^. These mutations are thought to stabilize an activated form of the FtsWI complex^5,9^, leading to increased rates of cell constriction^8^ via unrestrained PG synthesis.

If *fzlA* lies upstream of *ftsWI* in a PG synthesis pathway, then the hyperactive variants *ftsW**I\** and/or *ftsW\** may bypass the essentiality of *fzlA*. Accordingly, we found that *fzlA* could be readily deleted in either the *ftsW**I\** or *ftsW\** strain backgrounds (**Fig. 1A-B, Fig. S1A**). This is a particularly striking finding given that depletion of FzlA in a WT background completely inhibits division and induces cell filamentation and death^21^. Interestingly, a number of *ftsW**I*/ftsW\** Δ*fzlA* cells appeared to be “S”-shaped with the direction of curvature in future daughter cells facing opposite directions, as opposed to the characteristic “C”-shape of pre-divisional WT and *ftsW**I* Caulobacter* cells **(Fig. 1A**, asterisk, discussed further below). Assessment of fitness revealed that strains lacking *fzlA* displayed a slight reduction in viability by spot dilution, compared to the corresponding hyperactive PG synthase mutant strains (**Fig. 1C**), whereas growth rate was unaffected (**Fig. 1D**). In addition, *ftsW**I*/ftsW\** Δ*fzlA* cells displayed an increase in length (**Fig. 1E**), indicative of a residual division defect. Because *ftsW\** Δ*fzlA* cells are longer than *ftsW**I\** Δ*fzlA* cells, we conclude that *ftsW**I\** suppresses loss of *fzlA* better than the single mutant.

**Fig. 1:**
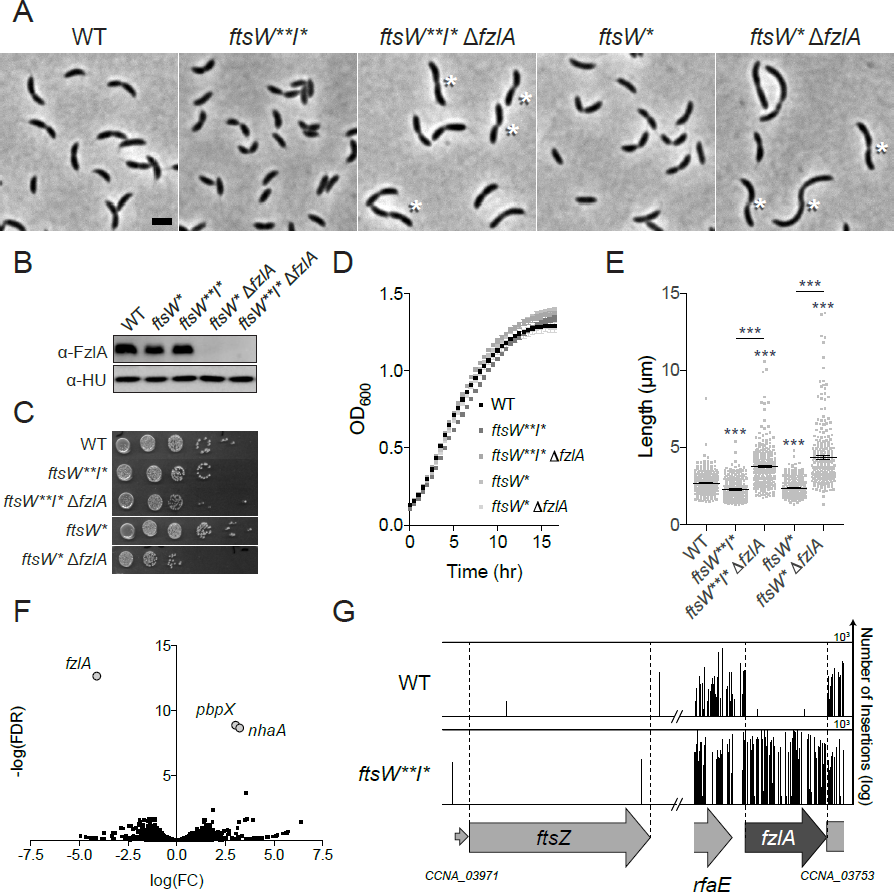
Hyperactive *ftsWI* mutants suppress loss of *fzlA*. A. Phase contrast microscopy images depicting WT *Caulobacter* and PG synthase hyperactive mutant cells with and without *fzlA*. White asterisks mark S-shaped cells. Scale bar = 2 µm. B. α-FzlA immunoblot (top) and α-HU immunoblot (bottom, loading control) of the indicated strains. C, D. Spot dilutions (ten-fold serial dilutions) (C) and growth curves (D) of the indicated strains. E. Lengths of unsynchronized cells from the indicated strains. Mean ± SEM shown. Kruskal-Wallis tests with Dunn’s post-test were performed to analyze differences compared to WT and the indicated strains: ^***^*P*≤0.001. From left to right, *n* = 254, 262, 261, 260, 258. F. Volcano plot of the negative log_10_ of the false discovery rate (-log(FDR)) vs. log_2_ of the fold change of each gene in WT vs. *ftsW**I\** strains determined by Tn-Seq analysis. G. Plot of transposon insertion frequency in essential division genes in WT (top) vs. *ftsW**I\** (bottom) cells determined by Tn-Seq analysis. Genetic loci are annotated below the plot. Number of reads is displayed on a logarithmic scale. Strain key (*Caulobacter crescentus*): WT (EG865 A-E; EG2366 F-G), *ftsW**I\** (EG1557), *ftsW**I\** Δ*fzlA* (EG2170), *ftsW\** (EG1556), *ftsW\** Δ*fzlA* (EG2166).

We also observed that *ftsW**I\** suppresses length, width, and fitness defects associated with slowly constricting *fzlA* point mutants *fzlA^NH2^* and *fzlA^NH3^* (**Fig. S2, Fig. S1B**), further indicating that hyperactivated *ftsWI* are dominant to, and likely downstream of *fzlA*. To determine the contribution of the FtsZ-FzlA interaction to activation of FtsWI, we assessed cell morphology, fitness, and cell length of *ftsW**I\** strains containing FzlA mutants with decreasing affinity for FtsZ^22^ (**Fig. S3, Fig. S1C**; FzlA > FzlA^NH2^ = FzlA^NH3^ > FzlA^NH1^; FzlA^NB2^, FzlA^NB1^ = no binding). We found that decreased affinity of FzlA towards FtsZ correlated with an increase in cell length (**Fig. S3E**), indicating that high-affinity binding to FtsZ is required for FzlA to signal to FtsWI.

### FzlA plays a specific and unique role in activating FtsWI

To assess the specificity of the *fzlA-ftsWI* genetic interaction and potentially identify additional components of this pathway, we performed comparative transposon sequencing (Tn-Seq) on WT and *ftsW**I\** strains. Surprisingly, *fzlA* was the only essential gene to become non-essential in the *ftsW**I\** background, with few insertions in WT but plentiful insertions in *ftsW**I\** cells (**Fig. 1F,G, Supplementary Table 1**). All other known essential division genes, e.g. *ftsZ* (**Fig. 1G**), had few transposon insertions in either background. These data indicate that *fzlA* is specific and unique in its essential role upstream of *ftsWI*. We suspect that other essential division proteins participate in this pathway as well, but that they play additional essential functions in divisome assembly or activity.

### FzlA contributes to efficient division in a hyperactive PG synthase background

Given that cells lacking *fzlA* in the hyperactive PG synthase backgrounds were elongated, we assessed constriction rate and division efficiency in these strains in more detail. Specifically, we performed time-lapse microscopy on *ftsW**I\** and *ftsW\** cells ± *fzlA* and tracked division in cells using MicrobeJ^22,23^ (**Fig. 2A, Supplementary Video 1**). Consistent with previous findings^8^, *ftsW**I\** and *ftsW\** cells constrict more quickly than WT (**Fig. 2B**). Intriguingly, the hyperactive PG synthase strains lacking *fzlA* constricted significantly more slowly than the corresponding strain with *fzlA* present, with constriction rates cut nearly in half (**Fig. 2B**). This suggests that hyperactivated FtsWI are not sufficient for efficient division and underscores the importance of FzlA in dictating constriction rate. As with cell length and fitness, *ftsW**I\** acted as a better suppressor to *fzlA* deletion, allowing for a faster constriction rate than did *ftsW\** (**Fig. 2B**).

**Fig. 2:**
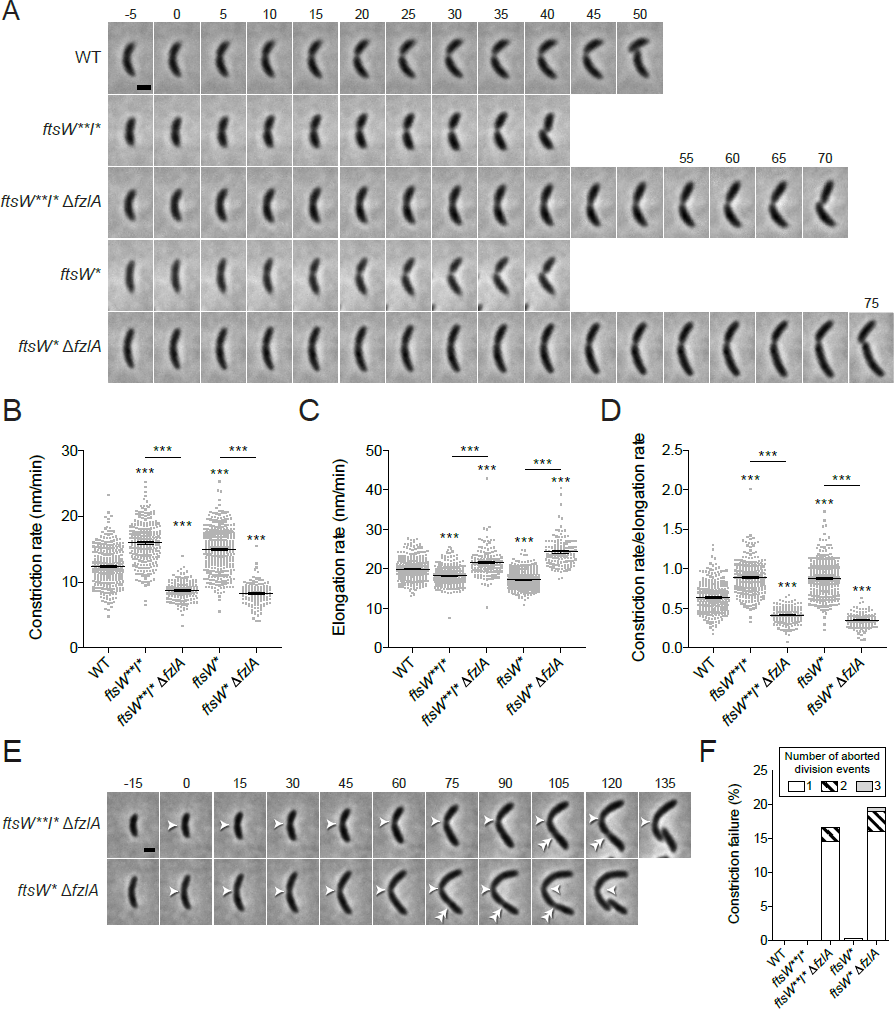
FzlA contributes to efficient division in a hyperactive PG synthase background. A. Phase contrast time-lapse microscopy images depicting constriction in WT or PG synthase hyperactive mutant cells with and without *fzlA*. Constriction starts at t=0 and concludes in the last frame upon cell separation. Time relative to constriction initiation (minutes) is indicated. Scale bar = 1 µm. B, C, D. Plots of constriction rate (B), total elongation rate (C), and ratio of constriction rate to total elongation rate (D) for a population of synchronized cells from each indicated strain, calculated from single cell microscopy data. Mean ± SEM shown. Kruskal-Wallis tests with Dunn’s post-test were performed to analyze differences compared to WT and the indicated strains: ^***^*P*≤0.001. From left to right, *n* = 324, 280, 161, 366, 139 (B) and 321, 280, 161, 363, 139 (C, D). E. Phase contrast microscopy images depicting constriction failure at the initial division site (single white arrowhead), then initiation and completion at a second site (double white arrowhead) in Δ*fzlA* cells. As in (A), constriction initiates at t=0 and concludes in the last frame. Scale bar = 1 µm. F. Plot of the constriction failure rate in cells in which constriction initiated, then failed at one division site and subsequently initiated and finished at another division site. “Number of aborted division events” refers to the number of times a cell abandoned division at distinct sites within the cell. The y-axis indicates the percentage of cells out of the whole population that displayed at least one such aborted division event. From left to right, *n* = 324, 280, 193, 368, 174. Strain key (*Caulobacter crescentus*): WT (EG865), *ftsW**I\** (EG1557), *ftsW**I\** Δ*fzlA* (EG2170), *ftsW\** (EG1556), *ftsW\** Δ*fzlA* (EG2166).

To ensure that changes in constriction rate were not due to global differences in PG synthesis, we determined elongation rates across strains (**Fig. 2C**), which enabled calculation of the ratio of constriction to elongation rate (**Fig. 2D**). We saw the same trend as for constriction rate itself, with *ftsWI\*\** and *ftsW\** mutant strains having higher ratios of constriction to elongation and loss of *fzlA* giving lower ratios (**Fig. 2D**). Interestingly, elongation rate was inversely correlated with constriction rate in all mutant strains (**Fig. 2C, Fig. 2D**), perhaps reflecting competition between the elongasome and divisome for PG precursor substrate^24^. Altogether, these data support the conclusion that alterations to the *ftsZ-fzlA-ftsWI* pathway specifically affect constriction, with FzlA increasing the constriction rate in both WT and hyperactive PG synthase mutant backgrounds.

While tracking division in *ftsW**I\** Δ*fzlA* and *ftsW\** Δ*fzlA* cells to measure constriction, we noticed that some cells initiated constriction at one location, then aborted division at that location before successfully dividing at a second (or third or fourth) site (**Fig. 2E, Supplementary Video 2**). We quantified the frequency of such constriction failure events and found that 16.6-19.5% of the hyperactive PG synthase cells lacking *fzlA* aborted division at one site before successfully dividing at another, compared to a 0-0.3% failure rate for WT or hyperactive PG synthase cells with *fzlA* present **(Fig. 2F)**. These data further demonstrate that *ftsW**I\** are not sufficient for efficient division, and that *fzlA* is required to ensure division processivity and efficiency.

### *fzlA* is required for maintenance of proper cell shape

As mentioned earlier, deletion of *fzlA* in the hyperactive PG synthase backgrounds impacted global cell morphology, with many pre-divisional cells appearing “S-shaped”. In order to more carefully assess this phenotype, we imaged cells by scanning electron microscopy (SEM). We saw a relatively high frequency of S-shaped *ftsW**I\** Δ*fzlA* cells, whereas most WT or *ftsW**I\** cells displayed the typical “C-shaped” morphology characteristic of *Caulobacter* (**Fig. 3A**). We quantified the frequency of S-shaped cells in a population of dividing cells by phase contrast microscopy to assess penetrance of this morphological phenotype. We extracted outlines of individual cells and performed principal component analysis using Celltool to isolate variance in cell shape to features referred to as shape modes^22,25^. Shape mode 3 captured the variation due to degree of S-versus C-shape and we set a cutoff such that cells with a standard deviation |sd| > 1 from the mean for this shape mode are considered S-shaped (**Fig. 3B,C**). Means and medians were similar for degree of S-shape across populations, with no significant difference for means, and a statistically significant but numerically small difference for medians. However, there was an obvious and significant difference in variance in degree of S-shape across populations (**Fig. 3B**), corresponding with a large difference in the number of cells found to be S-shaped in different strains. Over a quarter (26.9%) of dividing *ftsW**I\** Δ*fzlA* cells displayed an S-shaped morphology, compared to 2.4% of WT and 1.1% of *ftsW**I\** cells that are S-shaped (**Fig. 3D**).

**Fig. 3:**
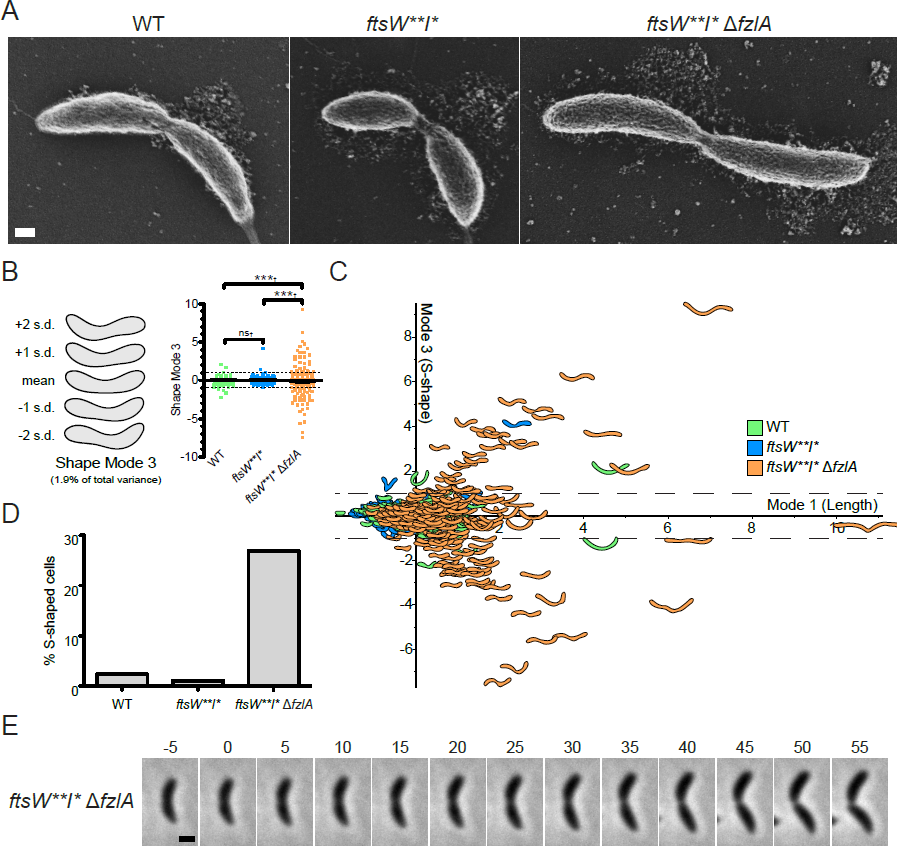
*fzlA* is required for global shape maintenance. A. SEM images of cells from the indicated strains. Scale bar = 200 nm. B. PCA of cell shape in a population of cells that have initiated constriction from the indicated strains. Shape mode 3 approximately captures degree of S-shape in cells. Mean cell contour ± 1 or 2 standard deviations (s.d.) is shown (left). Shape mode values for cells in each strain are plotted and mean ± SEM is shown (right). The dashed line drawn at s.d.=1 indicates the cutoff for S-shaped cells (cells with an |s.d.| ≥ 1 are considered to be S-shaped). A Brown-Forsythe Levene-type test (which is used in populations not assumed to be normally distributed) was performed to determine differences between population of population variances (†): *^ns^P>0.05*, ^***^*P*≤0.001. From left to right, *n* = 292, 279, 290. C. Plot of shape mode 3 (S-shape) vs. shape mode 1 (length) values for a cells that have initiated constriction from the indicated strains. The dashed lines drawn at |s.d.|=1 indicates the cutoff for S-shaped cells. D. Plot of percentage of S-shaped cells present in a population of cells that have initiated constriction from the indicated strains. Cells from (B) with an |s.d.| ≥ 1 are considered to be S-shaped. E. Phase contrast microscopy images depicting cell twisting during division in an *ftsW**I\** Δ*fzlA* cell. Constriction starts at t=0 minutes and concludes in the last frame upon cell separation. Scale bar = 1 µm. Strain key (*Caulobacter crescentus*): WT (EG865), *ftsW**I\** (EG1557), *ftsW**I\** Δ*fzlA* (EG2170).

To shed light on the origin of S-shape, we next asked at what point during growth do *ftsW**I\** Δ*fzlA* cells begin to adopt this morphology. Using time-lapse microscopy, we observed that *ftsW**I\** Δ*fzlA* cells were C-shaped at the beginning of the cell cycle and began to twist or rotate about the division plane after constriction initiated. S-shape only became apparent in the latter part of constriction, when daughters had rotated ~180° relative to each other (**Fig. 3E, Supplementary Video 3**). This finding suggests that the *fzlA-ftsWI* pathway determines geometry of PG insertion at the site of division in a manner that influences global cell morphology, normally constraining cells in their characteristic C-shape as constriction progresses. These results also indicate that our quantification method for S-shape likely underestimates the number of twisted *ftsW**I\** Δ*fzlA* cells, since S-shape is not obvious by phase contrast until the end of constriction, and we quantified cell shape at all stages of the constriction process.

Changes in division site shape and formation of S-shaped cells have been previously linked to aberrant localization of FtsZ and the elongation factor, MreB, respectively^19,26^. To determine if cell twisting might be facilitated by mislocalized division or elongation machineries, we visualized FtsZ and MreB localization in *ftsW**I\** Δ*fzlA* cells using inducible fluorescent fusions of these proteins. However, mNG-FtsZ and Venus-MreB localization in *ftsW**I\** Δ*fzlA* cells was comparable to *ftsW**I\** cells (**Fig. S4, S5**). Additionally, we visualized the localization of PG synthesis using the fluorescent D-amino acid HADA^27^, in order to assess whether cell twisting might be induced by mislocalized PG synthesis in spite of properly localized FtsZ and MreB. However, we did not detect any gross changes in HADA localization (**Fig. S6**). Together, these findings suggest that cell twisting is likely induced by a finer scale alteration of PG synthesis at the division site due to disruption of the *fzlA-ftsWI* pathway.

To determine whether global shape regulation depends on the FtsZ-FzlA interaction, we assessed S-shaped cell frequency in strains containing mutants of FzlA displaying decreasing affinities towards FtsZ. We observed that affinity of the mutant FzlA for FtsZ was inversely correlated with the frequency of S-shaped cells (**Fig. S7**), verifying that the FtsZ-FzlA interaction is important for maintaining proper morphology.

### The *ftsZ-fzlA-ftsWI* pathway contributes to resistance to PBP-targeting antibiotics

Because FzlA is important for regulation of PG synthesis in the context of determining constriction rate and cell shape, we hypothesized that it might also contribute to resistance to cell wall-targeting antibiotics. To test this, we challenged cells with antibiotics targeting PG synthetic processes and assessed resulting cell fitness. *ftsW**I*,* as has been previously shown^9^, displayed sensitivity to cephalexin (**Fig. 4A**), which inhibits FtsI and other penicillin-binding proteins in *Caulobacter* ^28,29^. Interestingly, deletion of *fzlA* in the *ftsW**I\** background exacerbated sensitivity to cephalexin (**Fig. 4A**). We found a similar trend upon treatment with mecillinam, which targets the elongation-specific PG synthase PBP2^30,31^, whereby the minimum inhibitory concentration (MIC) for *ftsW**I\** cells was decreased compared to WT, with deletion of *fzlA* further lowering the MIC (**Fig. 4B**). We also treated cells with the β-lactam ampicillin^32^ and with the cell wall targeting antibiotics vancomycin, which blocks transpeptidation by a distinct mechanism from β-lactams^33^, and fosfomycin, which inhibits cell wall synthesis by blocking PG precursor availability^34^ (**Fig. S8**). Neither hyperactivation of *ftsWI* nor loss of *fzlA* yielded a change in MIC in the presence of any of these antibiotics (**Fig. S8**). *fzlA* therefore supports robust cell wall synthesis in the presence of certain PG-targeting drugs, perhaps by compensating for inactivation of specific PBPs. To determine if the interaction between FtsZ and FzlA is important for maintaining cell wall integrity, we assessed sensitivity to cephalexin using the panel of *fzlA* mutants which display varying affinities towards FtsZ (**Fig. S9**). We found that mutants with decreased FzlA affinity towards FtsZ in fact became more sensitive to cephalexin (**Fig. S9**), demonstrating that the entire *ftsZ-fzlA-ftsWI* pathway is required for promoting cell wall integrity during antibiotic treatment.

**Fig. 4:**
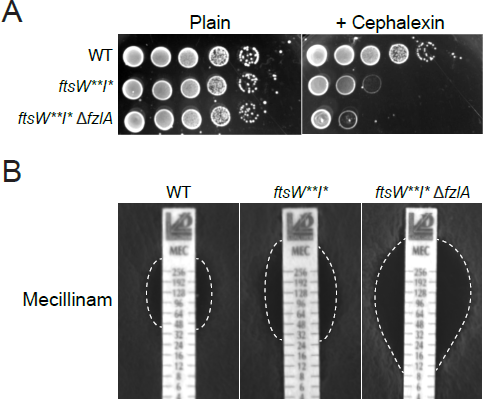
Loss of *fzlA* leads to increased cell wall antibiotic sensitivity. A. Spot dilutions (diluted ten-fold) of the indicated strains plated on PYE ± cephalexin (6 µg/ml). B. Plates of the indicated strains grown in the presence of mecillinam minimum inhibitory concentration (MIC) test strips, with antibiotic concentration decreasing from top to bottom. The zone of clearance is highlighted in white (dashed line). Strain key (*Caulobacter crescentus*): WT (EG865), *ftsW**I\** (EG1557), *ftsW**I\** Δ*fzlA* (EG2170).

Since *ftsW**I\** cells are more sensitive to perturbation of other PG synthetic activities even when *fzlA* is present, we asked if any normally non-essential division genes become more important for fitness in an *ftsW**I\** background, as they might help bolster resistance to assaults on PG synthesis. Examination of the *ftsW**I\** Tn-Seq data indicated that *pbpX* (encoding a bifunctional PG synthase that localizes to midcell)^35,36^, and to a lesser extent *ftsX* (encoding a cell separation factor)^37^ and *dipM* (encoding an envelope maintenance/cell separation factor)^37–40^, had fewer transposon insertions in an *ftsW**I\** background than in WT (**Fig. 1F, S10**). Because *ftsW**I\** cells have misregulated division site PG synthase activity, we suspect that FtsEX, DipM, and PbpX become important for ensuring robust PG synthesis during constriction and, later, efficient cell separation. Surprisingly, the normally non-essential *nhaA* locus, coding for a putative sodium-proton antiporter^41^, was also predicted by Tn-Seq to become essential in *ftsW**I\** cells (**Fig. 1F, Fig. S10B**). Disruption of *nhaA* in the presence of sucrose has been shown to arrest division^41^, suggesting *nhaA* may be important for division under certain conditions. It is unclear why it also becomes important upon PG synthesis mis-regulation, but its role in osmoregulation may contribute to its apparent synthetic lethality with *ftsW**I\**.

### The *fzlA-ftsW* genetic interaction is conserved in diverse α-proteobacteria

FzlA homologs are encoded in nearly all sequenced α-proteobacterial genomes, but not outside this group. To assess the conservation of FzlA’s role in regulating PG synthesis, we sought to characterize the genetic interaction between *fzlA* and PG synthases in another α-proteobacterium, *Agrobacterium tumefaciens*. *A. tumefaciens* and *Caulobacter* display disparate growth patterns driven by distinct machineries during elongation, with *A. tumefaciens* exhibiting polar elongation and *Caulobacter* elongating primarily at midcell and through dispersed growth ^27,42–44^. However, the components of the division machinery, including FtsZ, FzlA, and FtsWI, are largely conserved. To test if the genetic interaction between *fzlA* and *ftsW* is conserved, we made an IPTG-dependent FzlA depletion construct in a WT background or in a background with a single hyperactivating mutation in *A. tumefaciens ftsW* (F137L, the equivalent of *Caulobacter* FtsW F145L) at the *ftsW* locus. Depletion of FzlA in a WT background resulted in reduced viability, FtsW depletion in *A. tumefaciens*^18^. Importantly, we found that the decrease in viability division arrest, and ectopic pole formation at midcell (**Fig. 5, Fig. S11**), reminiscent of and morphology defects associated with depletion of FzlA were rescued by *ftsW F137L* (**Fig. 5, Fig. S11**). These data indicate that FzlA’s essential role in regulating division-specific PG synthesis is conserved in another α-proteobacterium and further highlight the importance of FzlA as a key regulator of constriction and cell morphology.

**Fig. 5:**
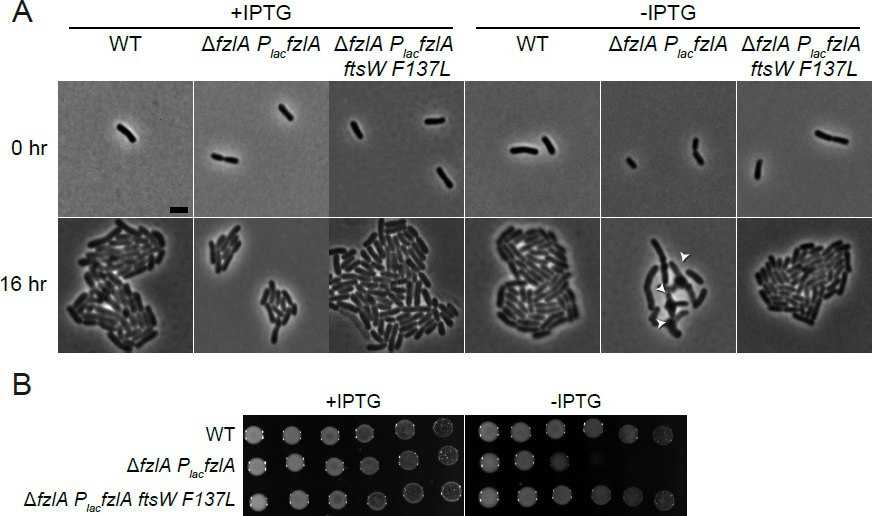
The ability of hyperactive *ftsW* to suppress loss of *fzlA* is conserved. A. Phase contrast microscopy images depicting PG synthase hyperactive mutant cells ± FzlA in *Agrobacterium tumefaciens*. FzlA was induced where indicated with IPTG and depleted where indicated upon removal of IPTG, then grown for 16 hours on agarose pads. White arrowheads mark ectopic poles at midcell. Scale bar = 2 µm. B. Spot dilutions (ten-fold serial dilutions) of the indicated strains grown in the presence or absence of IPTG to control *fzlA* expression, in *A. tumefaciens*. Strain key (*A. tumefaciens*): WT (PBA44)^48^; Δ*fzlA P_lac_fzlA* (PBA199); Δ*fzlA P_lac_fzlA ftsWF137L* (PBA232).

## Discussion

Here we have described a conserved PG synthesis activation pathway in which FtsZ and FzlA signal through FtsWI to regulate wall synthesis during division in α-proteobacteria (**Fig. 6, left panel**). Specifically, the FtsZ-FzlA-FtsWI pathway determines geometry of cell wall insertion at the site of division, sets the constriction rate, and promotes cell wall integrity (**Fig. 6, left panel**). FtsW**I* can still receive input from FzlA which, in combination with their intrinsic hyperactivity, leads to shorter, faster-constricting cells with sensitivity to cell wall antibiotics (**Fig. 6, middle panel**). In the absence of *fzlA*, *ftsW**I\** cells lose critical regulation of PG synthesis, leading to twisting during division, slower constriction, and increased sensitivity to cell wall antibiotics. We establish FzlA as a key intermediary in signaling from FtsZ to FtsWI and demonstrate that this division-specific SEDS-PBP pair require activation for normal division. Notably, our observations indicate that FtsWI activity is regulated in multiple ways, likely including input into their catalytic rates and modulation of the fine-scale geometry by which they insert new PG for constriction.

**Fig. 6:**
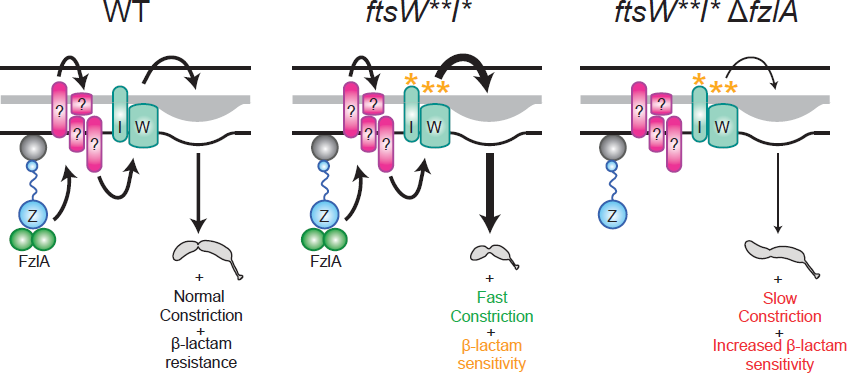
FzlA is required for activation of FtsWI and regulates the geometry of PG insertion. FzlA is required for activation of FtsWI and likely signals through unidentified intermediate factor(s) in a manner dependent on interaction with FtsZ to effect normal cell shape, normal constriction rate, and antibiotic resistance (left). FtsW**I* can still receive input from FzlA, which in combination with its own hyperactivity, leads to faster constriction and antibiotic sensitivity likely associated with positive misregulation of PG insertion (middle). FtsW**I* can function in the absence of FzlA, but with misregulated activity, leading to twisting during constriction, slower constriction speed, and increased antibiotic sensitivity (right).

Our findings provide the foundation for further mechanistic investigation into this pathway and raise a number of questions. For one, the nature and timing of the activation signal(s) are still unknown: is there a signal always emanating from FtsZ-FzlA that induces constriction as soon as FtsW arrives, or is constriction triggered by a discrete cellular event, such as clearance of the chromosomal termini or the arrival of a sparkplug factor that jumpstarts FtsWI activityŒ Additionally, it is unclear why hyperactivation of *ftsWI* together or *ftsW* alone can rescue loss of *fzlA* – are TPase and GTase activities impacted similarly, or does one predominate in regulating constriction rateŒ A recent described here in fact hyperactivates FtsW. Finally, we have no evidence that FzlA and study demonstrated that FtsW is activated by FtsI^7^, so it is possible that the *ftsI\** mutation FtsWI directly interact, and suspect that other intermediary factor(s) transduce the activation signal from FzlA to FtsWI.

Our model advances the idea that regulation of SEDS-PBP pairs for growth and division is conserved at numerous levels. The finding that FzlA governs division-specific PG synthesis in both *Caulobacter* and *A. tumefaciens* argues that α-proteobacteria use FzlA as a conserved and dedicated FtsWI activator. FzlA is absent outside of this clade, however, so we propose that other divisome components serve as FtsWI activators in other organisms. More broadly, our findings expand the paradigm for PG synthesis by SEDS-PBP PG synthase pairs in bacteria and provide evidence that the requirement for PG synthase activation is conserved. Elongation is facilitated by the coordination of the SEDS family GTase RodA and the monofunctional TPase PBP2, orthologs to FtsW and FtsI, respectively^5^. The proposed model for elongation activation, as described for *E. coli*, holds that these PG synthases are activated by another protein, MreC, forming an activated complex that in turn regulates assembly and directional motion of the polymerizing scaffold MreB^5^. In this system, hyperactivating mutations in RodA or PBP2 allow for bypass of the activator, MreC, similar to our finding that FtsW**I* can bypass the activator FzlA. Our data provide experimental support for the proposal that the requirement for activation of the SEDS-PBP pair of PG synthases is generally conserved for elongation and division.

There are prominent differences between the models for elongasome and divisome activation, however. The elongasome appears, in essence, to be a stripped down and when either RodA or PBP2 is hyperactivated through mutation, all elongasome version of the divisome^45^: the elongasome contains fewer proteins than the divisome^45^, components except MreB and the PG synthases are rendered dispensable^5^. This would suggest that for elongation, the cell needs an activated SEDS-PBP pair and a spatial regulator to orient their motion^5^. Conversely, hyperactive FtsWI in *Caulobacter* only allows for disruption of FzlA, with the rest of the divisome remaining essential. This may be because division is a more complex process than elongation, requiring invagination and fission of all layers of the envelope in coordination with DNA segregation and cell cycle progression. This complexity necessitates functions in addition to PG synthesis and remodeling provided by components of the divisome. PG synthesis during division likely requires more regulation, as well. Whereas PG synthesis during elongation comprises insertion of new PG in the same plane as old cell wall material, PG synthesis during constriction requires a lasting, directional change to shape the new cell poles. So while there are key similarities in the paradigm of PG synthase activation, regulation of division likely requires a more complicated network of inputs to manage the additional outputs and constraints discussed above. In summation, this work provides evidence that the requirement for SEDS-PBP activation is conserved across multiple modes of PG synthesis, which has broad implications for determining the speed of division, cell shape, and cell wall robustness.

## Methods

### Strains, growth conditions, and growth determination

*Caulobacter crescentus* strains were derived from the NA1000 WT strain. Unless otherwise indicated, *Caulobacter* colonies were isolated from solid 1.5% agar peptone yeast extract (PYE) plates grown at 30 °C and cells were grown in liquid culture in PYE shaking at 30 °C. Where indicated, *Caulobacter* cells were treated with 6 µg/ml of cephalexin. Antibiotic MIC analysis was performed using antibiotic test strips (Liofilchem), which include a concentration gradient of 0.016-256 mg/L for all antibiotics tested. Where indicated, cells were treated with 0.3% xylose to drive inducible gene expression. Spot dilutions for *Caulobacter* were performed by serially diluting cells at the indicated fraction (1/10 or 1/2), before plating. Growth rates were obtained by measuring optical density at 600 nm (OD_600_) values of cells every 30 minutes. *Caulobacter* cell synchrony was performed as previously described^12^. Briefly, log phase cells were washed with M2 salts (6.1 mM Na_2_HPO_4_, 3.9 mM KH_2_PO_4,_ 9.3 mM NH_4_Cl)^46^, resuspended in 1:1 M2:Percoll, then centrifuged at 11,200 x *g*. The swarmer band was isolated, and cells were subsequently washed twice in M2, then resuspended in PYE.

*A. tumefaciens* were grown in *A. tumefaciens* glucose and (NH_4_)_2_SO_4_ (ATGN) minimal medium^47^, with 0.5% glucose at 28°C. *E. coli* strains were grown in LB medium at 37°C. IPTG was added at a concentration of 1 mM when necessary. For *A. tumefaciens* spot dilutions, cells were grown overnight in ATGN minimal medium in the presence of IPTG at 1 mM concentration, washed, then pre-depleted of IPTG for 16 hours where indicated. Cells were then serially diluted (ten-fold) and spotted on ATGN minimal medium with the presence or absence of IPTG. To make the Δ*fzlA P_lac_fzlA* strain (PBA199), first, a mini-Tn7 vector containing IPTG inducible *fzlA*, along with the pTNS3 helper plasmid, were introduced into D*tetRA* a-*att*Tn7 cells (PBA44) via electroporation as previously described^48^. Deletion of *fzlA* (for PBA199) and allelic exchange of *ftsW* (for PBA232) were subsequently performed by transferring the corresponding suicide vector to *A. tumefaciens* via conjugation with *E. coli* S17.

Plasmids (**Supplementary Table 2**) and strains (**Supplementary Table 3**) used in this study can be found in the supplementary information. Plasmids were created using standard molecular cloning procedures including PCR, restriction digestion, and ligation. Mutagenesis of *ftsW* for *A. tumefaciens* was performed using a QuikChange Lightning Multi Site­Directed Mutagenesis Kit (Agilent Genomics), with primers designed using Agilent’s QuikChange Primer Design Program, as previously described^22^. pEG1345 was constructed using an NEBuilder HiFi DNA Assembly Cloning Kit (NEB).

### Light microscopy imaging and analysis

Images of log phase *Caulobacter* cells were obtained using either phase contrast microscopy, with cells grown on either 1% agarose PYE pads or 1% agarose dH_2_O pads, or, when indicated, fluorescence microscopy, with cells grown on 1% agarose dH_2_O pads. For fluorescence microscopy, mNG-FtsZ expression was induced for 1 hour with xylose then imaged through the GFP filter and venus-MreB expression was induced for 2 hours with xylose then imaged through the YFP filter. For determination of PG incorporation localization, cells were pulsed with 0.82 mM HADA for 5 minutes, washed twice with PBS, then visualized through the DAPI filter. For time-lapse imaging, as previously described^22^, synchronized cells were placed on 1% agarose PYE pads and imaged using phase contrast microscopy at room temperature (RT), with images being acquired at 5 minute intervals at 100x. Imaging of *Caulobacter* cells was performed using a Nikon Eclipse Ti inverted microscope with a Nikon Plan Fluor × 100 (numeric aperture 1.30) oil Ph3 objective and Photometrics CoolSNAP HQ^2^ cooled CCD (charge­coupled device) camera^22^. For *A. tumefaciens* phase contrast microscopy, exponentially growing cells were spotted on 1% agarose ATGN pads as previously described^49^, then imaged. For *A. tumefaciens* time-lapse microscopy, images were collected every ten minutes. Microscopy of *A. tumefaciens* cells was performed with an inverted Nikon Eclipse TiE with a QImaging Rolera em-c^2^ 1K EMCCD camera and Nikon Elements Imaging Software.

For determination of dimensions of log phase cells, cell length and width were measured using MicrobeJ software, similar to as previously described^22^. Constriction rate and elongation rate were also determined using MicrobeJ^23^. Briefly, MicrobeJ software allowed for tracking of cells imaged by time-lapse microscopy throughout the division process, with automatic detection of constriction initiation and manual determination of cell separation. Cell length was found for cells at each time point, cell width was found at the site of constriction, and constriction time was calculated by multiplying the number of frames in which constriction was detected by 5 (since images were acquired every 5 minutes), allowing for calculation of constriction and elongation rates. Constriction failure rate was determined by counting the number of cells which initiated constriction at one division site, failed, then ultimately divided at a separate site. Prism was used for graphing and statistical analysis of calculated terms.

### Cell shape analysis

For cell shape analysis, binary masks of phase contrast images of log phase cells were inputted into Celltool^25^, allowing for creation of cell contours, similar to as previously described^22^. Following alignment of cell contours (not allowing for reflection), a model of cell shape was created. The shape modes of interest were either plotted as histograms displaying the cell shape across two dimensions, or as single data points. R software was used to perform statistical analyses to compare population variances in shape modes across strains. Prism was used for graphing calculated terms.

### Scanning electron microscopy sample preparation and imaging

For SEM, log phase cells were incubated on poly-lysine (1:10) coated glass cover slip for 15 minutes, then fixed for 1 hour using fixation buffer (1% glutaraldehyde, 0.02 M cacodylate, and 3 mM MgCl_2_). Cells were gradually dried by washing 3 times with wash buffer (3% sucrose, 0.02 M cacodylate, and 3 mM MgCl_2_), twice with dH_2_O, once each with 30%, 50%, 70%, 90%, 100% ethanol, once with 1:1 ethanol:hexamethyldisiloxane (HMDS), and once with HMDS at 5 minute intervals each, before desiccation overnight. Cover slips were mounted, then coated with a 15 nm gold palladium sputter coat. Samples were then imaged with a LEO/Zeiss Field-emission SEM.

### Immunoblot analysis

Immunoblot analysis was performed similar to as previously described^22^, using a 1:5,000 – 1:6,666 dilution of α-FzlA primary antibody^21^, a 1:50,000 dilution of α-HU primary antibody^50^, and 1:10,000 of HRP-labeled α-rabbit secondary antibody (PerkinElmer) on nitrocellulose membranes. Chemiluminescent substrate (PerkinElmer) was added to facilitate protein visualization via an Amersham Imager 600 RGB gel and membrane imager (GE).

### Transposon library preparation, sequencing, and analysis

Wild type *Caulobacter crescentus* NA1000 (EG2366) or *ftsW**I\** triple mutant (EG1557) cells were grown in a large culture (1 liter) to mid-log (0.4-0.6), washed of excess Mg^2+^ with 10% glycerol, and mutagenized with the Ez-Tn5 <Kan-2> transposome (Epicentre). Cells recovered by shaking at 30 °C for 90 minutes, then plated on kanamycin containing plates for 3 days at 30 °C in order to yield roughly 100-500 colonies per plate. Libraries were grown at 30 °C and comprised ~100,000-200,000 colonies each. Mutants were pooled into one library by scraping colonies from the surface of the agar and added into ~25-40 mL PYE. Pooled libraries were shaken to yield a homogenous slurry and sterile glycerol was added to 20%. Libraries were then frozen in liquid nitrogen and stored at −80 °C. Two libraries of each genetic background were prepared individually and compared as biological replicates.

Genomic DNA was extracted from one aliquot of each library using DNeasy Blood and Tissue Kit (Qiagen). Tn-Seq libraries were prepared for Illumina Next-Generation sequencing through sequential PCR amplifications using arbitrary hexamer primers and Tn5-specific primer facing outward for the first round, and indexing primer sets that include unique molecular identifier to filter artifacts arising from PCR duplicates for the second round. Libraries were then pooled and sequenced at the University of Massachusetts Amherst Genomics Core Facility on the NextSeq 550 (Illumina).

For analyses, reads were demultiplexed by index, then each sample Tn-Seq library was concatenated and clipped of the unique molecular identifier linker from the second PCR using Je^51^ and the following command:

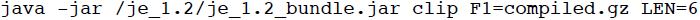

Clipped reads were then mapped back to the *Caulobacter* NA1000 genome (NCBI Reference Sequence: NC_011916.1) using BWA^52^ and sorted using Samtools^53^:

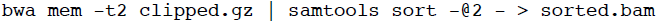

Duplicate reads were removed using Je^51^ and indexed with Samtools^53^ using the following command:

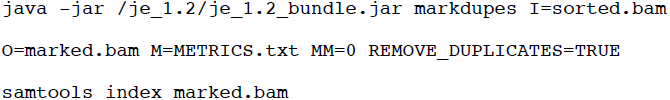

5’ sites of inserted transposons from each library were converted into .wig files containing counts per position and viewed using Integrative Genomics Viewer^54,55^. Coverage and insertion frequency using a bedfile containing all open reading frames from NC_011916.1 with the outer 20% of each gene removed were determined using BEDTools^56^ and the following commands:

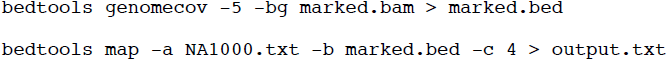

Comparison of transposon insertions was performed using the edgeR package in the Bioconductor suite^57,58^ using a quasi-likelihood F-test (glmQLFit) to determine the false discovery rate adjusted p-values reported here.

## Acknowledgements

We would like to thank Josh Modell and Mike Laub for providing strains; the Manley lab, especially Ambroise Lambert, and members of the Xiao and Goley labs for helpful discussions; the Goley lab for feedback on the manuscript; Anant Bhargava and Adrien Ducret for help with time-lapse analysis; Mike Delannoy, Barbara Smith, and Selam Woldemeskel for developing the SEM protocol, and Mike and Barbara for training on SEM equipment; and Brandon King for providing support with experiments. This work funded in part by the National Institutes of Health, National Institute of General Medical Sciences through R01GM108640 (EDG and PJL), T32GM007445 (training grant support of PJL), R01GM111706 (PC and RZ), R25GM056901(training support of GS-C), and T32GM08515 (training grant support of RZ). PB and MH were supported by the National Science Foundation, IOS1557806.

## Author Information

PJL, CRM, AKD, PB, MH, GS-C, PC, RZ, and EDG planned the experiments, PJL, CRM, AKD, MH, GS-C, and RZ performed the experiments, PJL, CRM, AKD, MH, and EDG wrote the manuscript, and PJL, CRM, AKD, PB, MH, GS-C, PC, and EDG edited the manuscript.

## Supplementary Information

**Supplementary Figure Legends**

**Fig. S1:**
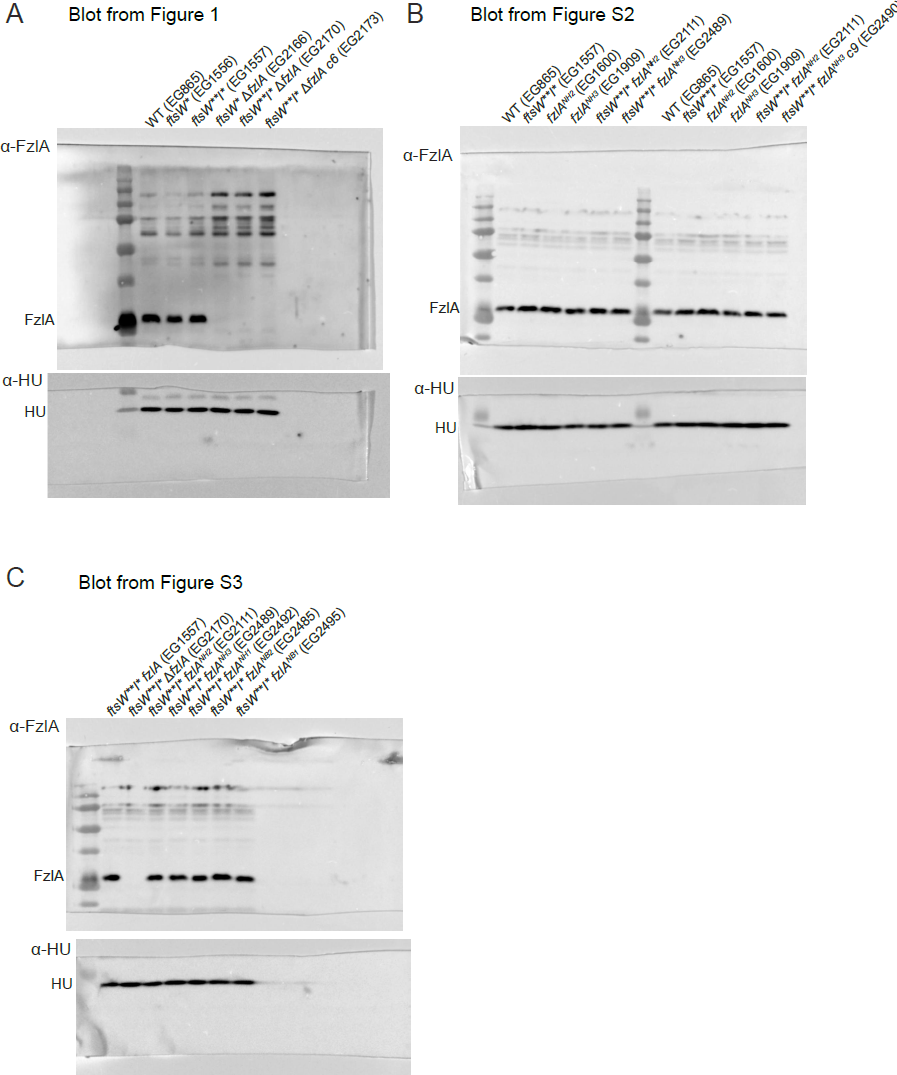
Uncropped Immunoblots. A-C. Uncropped α-FzlA immunoblots (top) and α-HU immunoblots (bottom, loading control) of the indicated strains from figure 1 (A), figure S2 (B, left half), and figure S3 (C).

**Fig. S2:**
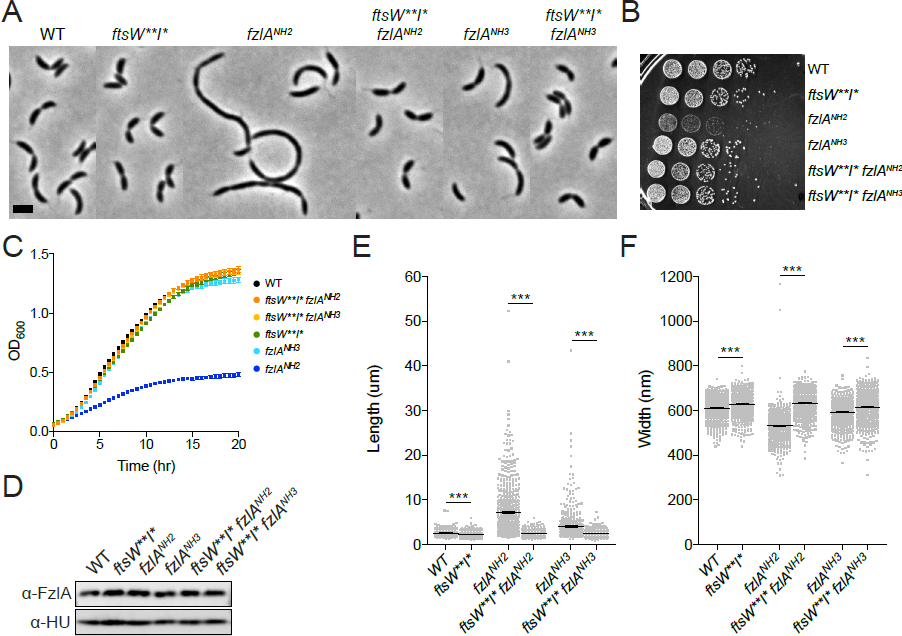
*ftsW**I\** rescue the fitness/morphological defects of two *fzlA* mutants. A. Phase contrast microscopy images depicting cells of the indicated strains. Scale bar = 2 µm. B, C. Spot dilutions (diluted ten-fold) (B) and growth curves (C) of the indicated strains. D. α-FzlA immunoblot (top) and α-HU immunoblot (bottom, loading control) of the indicated strains. E, F. Lengths (E) and widths (F) of unsynchronized cells from the indicated strains. Mean ± SEM shown. Kruskal-Wallis tests with Dunn’s post-test were performed to analyze differences compared to the indicated strains: ^***^*P*≤0.001. From left to right, *n* = 674, 609, 606, 653, 618, 645. Strain key (*Caulobacter crescentus*): WT (EG865), *ftsW**I* fzlA* (EG1557), *fzlA^NH2^* (EG1600), *ftsW**I* fzlA^NH2^* (EG2111), *fzlA^NH3^* (EG1909), *ftsW**I* fzlA^NH3^* (EG2489).

**Fig. S3:**
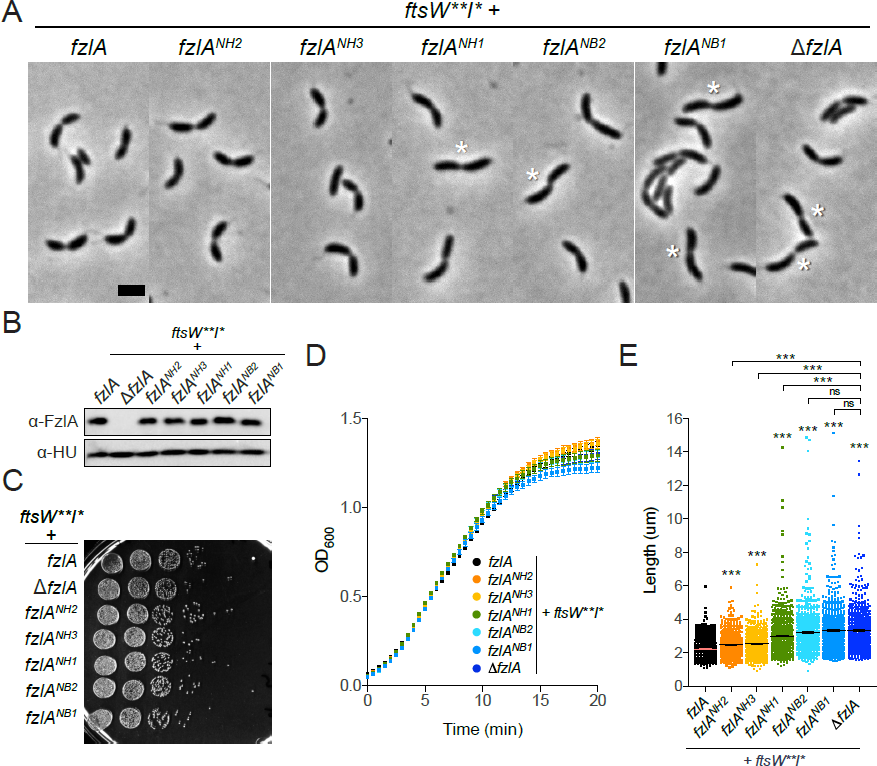
In the presence of hyperactive PG synthases, the interaction between FtsZ and FzlA determines division efficiency, but not growth rate or viability. A. Phase contrast microscopy images depicting cells of the indicated strains. White asterisks mark S-shaped cells. Scale bar = 2 µm. B. α-FzlA immunoblot (top) and α-HU immunoblot (bottom, loading control) of the indicated strains. C, D. Spot dilutions (diluted ten-fold) (C) and growth curves (D) of the indicated strains. E. Lengths of unsynchronized cells from the indicated strains. Mean ± SEM shown. Kruskal-Wallis tests with Dunn’s post-test were performed to analyze differences compared to WT and the indicated strains: ^*ns*^P>0.05, ^***^*P*≤0.001. From left to right, *n* = 609, 653, 645, 688, 674, 729, 612. Strain key (*Caulobacter crescentus*): *ftsW**I* fzlA* (EG1557), *ftsW**I* fzlA^NH2^* (EG2111), *ftsW**I* fzlA^NH3^* (EG2489), *ftsW**I* fzlA^NH1^* (EG2492), *ftsW**I* fzlA^NB2^* (EG2485), *ftsW**I* fzlA^NB1^* (EG2495), *ftsW**I\** Δ*fzlA* (EG2170).

**Fig. S4:**
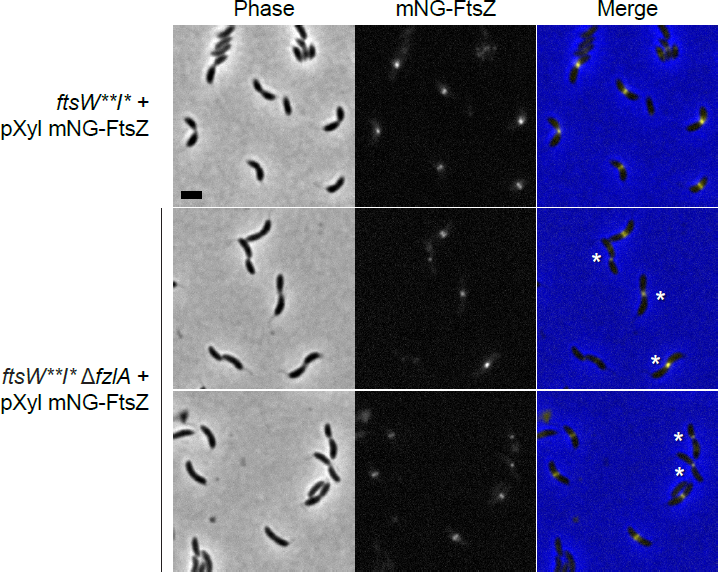
FtsZ localization is unaffected in *ftsW**I\** Δ*fzlA* cells. Phase contrast, fluorescence, and merged microscopy images depicting mNG-FtsZ localization in cells of the indicated strains in the presence of inducer (xylose). White asterisks mark S-shaped cells. Scale bar = 2 µm. Strain key (*Caulobacter crescentus*): *ftsW**I\** + *pXyl mNG-ftsZ* (EG2157), *ftsW**I\** Δ*fzlA* + *pXyl mNG-ftsZ* (EG2326).

**Fig. S5:**
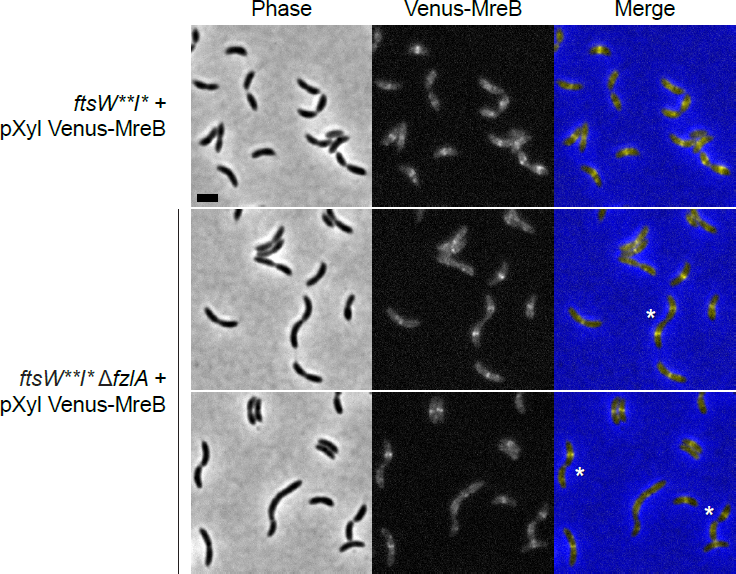
MreB localization is unaffected in *ftsW**I\** Δ*fzlA* cells. Phase contrast, fluorescence, and merged microscopy images depicting Venus-MreB localization in cells of the indicated strains in the presence of inducer (xylose). White asterisks mark S-shaped cells. Scale bar = 2 µm. Strain key (*Caulobacter crescentus*): *ftsW**I\** + *pXyl Venus-mreB* (EG2377), *ftsW**I\** Δ*fzlA* + *pXyl Venus-mreB* (EG2378).

**Fig. S6:**
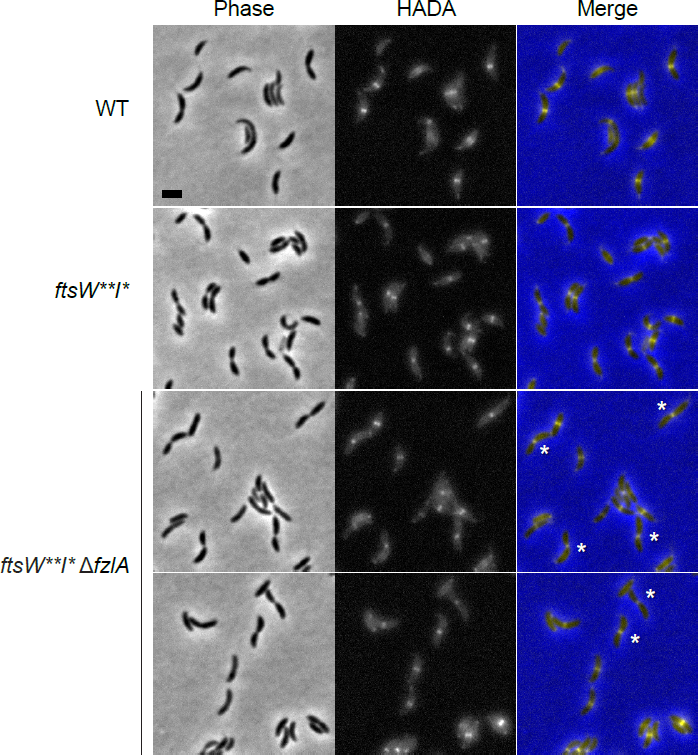
The localization of PG synthesis is unaffected in *ftsW**I\** Δ*fzlA* cells. Phase contrast, fluorescence, and merged microscopy images depicting HADA localization in cells of the indicated strains after a 5 minute HADA pulse. White asterisks mark S-shaped cells. Scale bar = 2 µm. Strain key (*Caulobacter crescentus*): WT (EG865), *ftsW**I\** (EG1557), *ftsW**I\** Δ*fzlA* (EG2170).

**Fig. S7:**
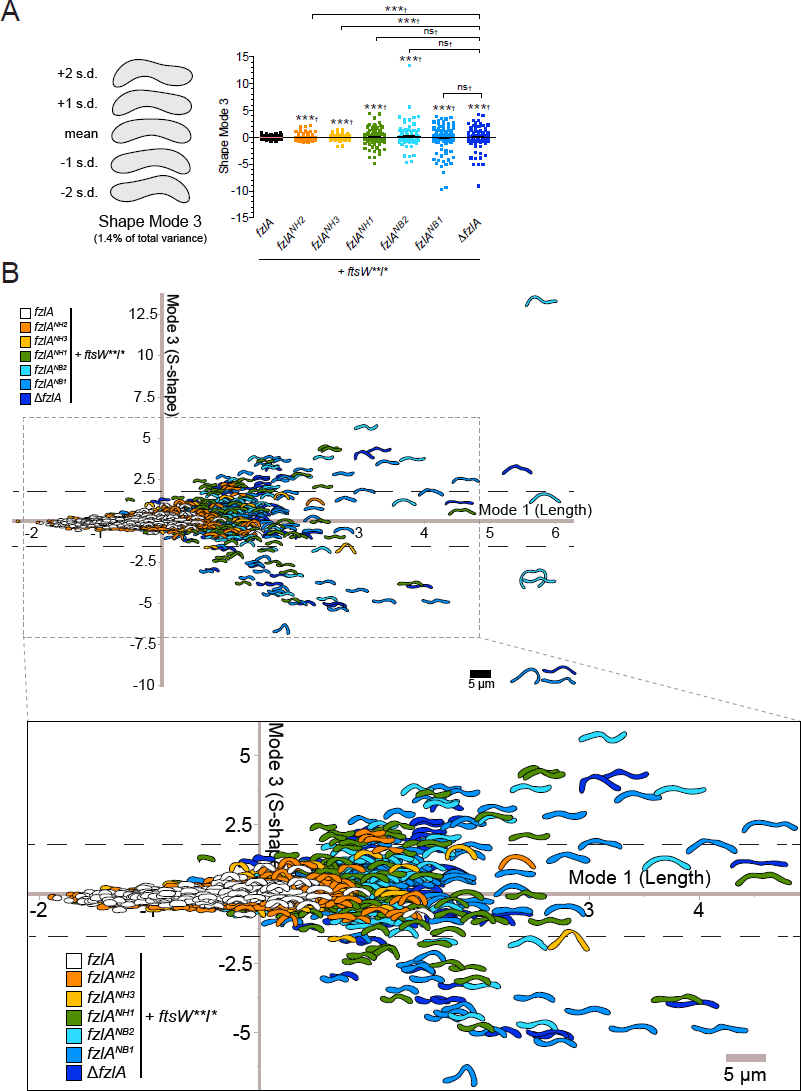
Interaction of FtsZ with FzlA is necessary for proper division site shape maintenance. A. PCA of cell shape in a population of unsynchronized cells (not all cells are necessarily actively constricting) from the indicated strains. Shape mode 3 approximately captures degree of S-shape in cells. Mean cell contour ± 1 or 2 standard deviations (s.d.) is shown (left). Shape mode values for cells in each strain are plotted and mean ± SEM is shown (right). A Brown-Forsythe Levene-type test (which is used in populations not assumed to be normally distributed) was performed to determine differences between population variances (†): ^*ns*^P>0.05, ^***^*P*≤0.001. From left to right, *n* = 269, 375, 218, 250, 177, 289, 211. B. Plot of shape mode 3 (degree of S-shape) vs. shape mode 1 (length) values in a population of unsynchronized cells (not all cells are necessarily actively constricting) from the indicated strains. The dashed lines qualitatively demark the boundary between cells that appear to be S-shaped and those that display normal curvature. Inset presents a zoomed in view of the highlighted region of interest. Strain key (*Caulobacter crescentus*): *ftsW**I* fzlA* (EG1557), *ftsW**I* fzlA^NH2^* (EG2111), *ftsW**I* fzlA^NH3^* (EG2489), *ftsW**I* fzlA^NH1^* (EG2492), *ftsW**I* fzlA^NB2^* (EG2485), *ftsW**I* fzlA^NB1^* (EG2495), *ftsW**I\** Δ*fzlA* (EG2170).

**Fig. S8:**
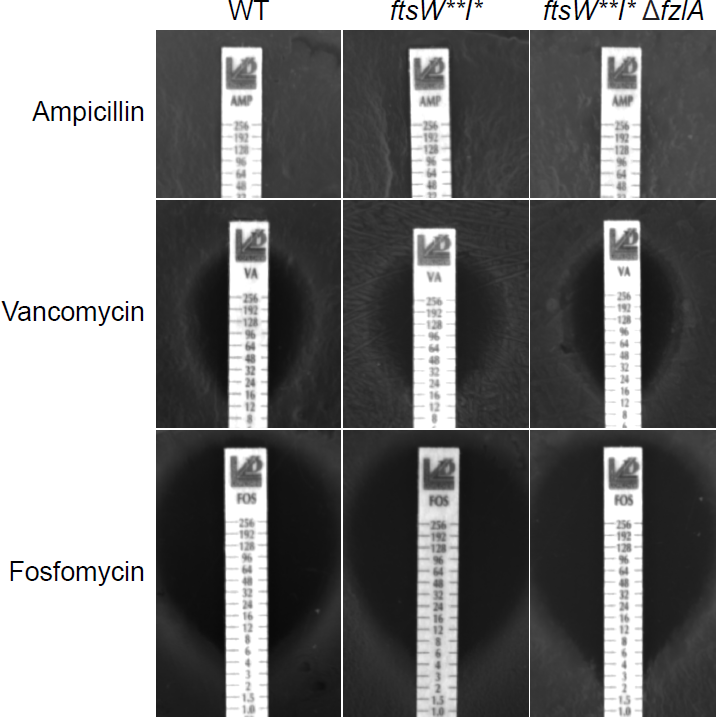
Loss of *fzlA* does not confer increased sensitivity to various classes of antibiotics. Plates of the indicated strains grown in the presence of antibiotic minimum inhibitory concentration (MIC) test strips, with antibiotic concentration decreasing from top to bottom. Strain key (*Caulobacter crescentus*): WT (EG865), *ftsW**I\** (EG1557), *ftsW**I\** Δ*fzlA* (EG2170).

**Fig. S9:**
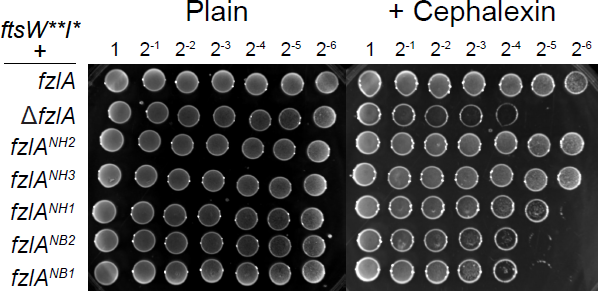
Interaction of FtsZ with FzlA is necessary for increased resistance to cell wall antibiotics. Spot dilutions (diluted two-fold) of the indicated strains plated on PYE ± cephalexin (6 µg/ml). Strain key (*Caulobacter crescentus*): *ftsW**I* fzlA* (EG1557), *ftsW**I* fzlA^NH2^* (EG2111), *ftsW**I* fzlA^NH3^* (EG2489), *ftsW**I* fzlA^NH1^* (EG2492), *ftsW**I* fzlA^NB2^* (EG2485), *ftsW**I* fzlA^NB1^* (EG2495), *ftsW**I\** Δ*fzlA* (EG2170), *fzlA^NH2^* (EG1600), *fzlA^NH3^* (EG1909).

**Fig. S10:**
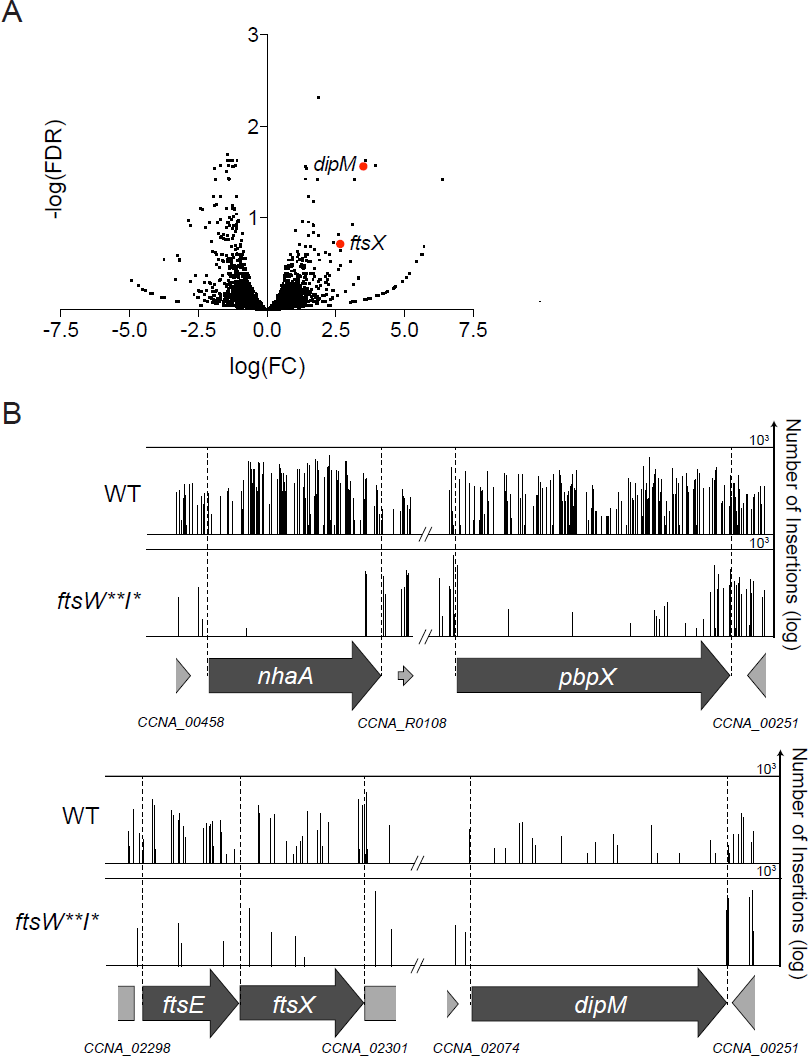
Multiple non-essential division genes become essential in a hyperactive PG synthase background. A. Volcano plot of the negative log_10_ of the false discovery rate (-log(FDR)) vs. log_2_ of the fold change of each gene in WT vs. *ftsW**I\** strains determined by Tn-Seq analysis. This is a zoomed in and cropped view of the volcano plot from Figure 1F. B. Plot of transposon insertion frequency in essential division genes in WT (top) vs. *ftsW**I\** (bottom) cells. Genetic loci are annotated below the plot. Number of reads is displayed on a logarithmic scale. Strain key (*Caulobacter crescentus*): WT (EG2366), *ftsW**I\** (EG1557)

**Fig. S11:**
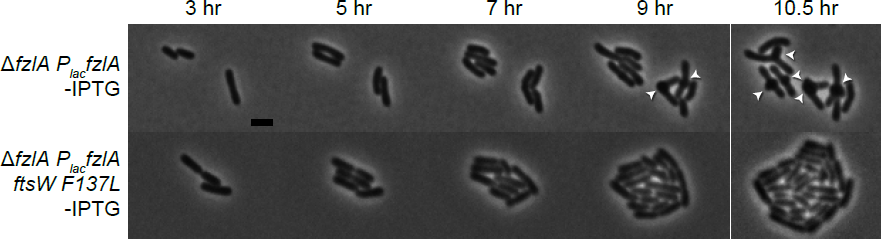
Time-lapse showing hyperactive *ftsW* suppresses loss of *fzlA* in *A. tumafaciens*. Phase contrast time-lapse microscopy images depicting WT and PG synthase hyperactive mutant cells depleted of FzlA over time. White arrowheads mark ectopic poles at midcell. Scale bar = 2 µm. Strain key (*Agrobacterium tumefaciens*): Δ*fzlA P_lac_fzlA* (PBA199); Δ*fzlA P_lac_fzlA ftsWF137L* (PBA232).

## Supplementary Video Legends

**Supplementary Video 1:**

Phase contrast time-lapse microscopy movies depicting division in WT _or_ PG synthase hyperactive mutant cells with and without *fzlA*. As indicated, 5 minutes elapse between frames. Video playback is 10 frames per second.

Strain key (*Caulobacter crescentus*): WT (EG865), *ftsW**I\** (EG1557), *ftsW**I\** Δ*fzlA* (EG2170), *ftsW\** (EG1556), *ftsW\** Δ*fzlA* (EG2166).

**Supplementary Video 2:**

Phase contrast time-lapse microscopy movies depicting examples of constriction failure at the initial division site then initiation and completion at a subsequent site in Δ*fzlA* cells. As indicated, 5 minutes elapse between frames. Video playback is 10 frames per second.

Strain key (*Caulobacter crescentus*): *ftsW**I\** Δ*fzlA* (EG2170), *ftsW\** Δ*fzlA* (EG2166).

**Supplementary Video 3:**

Phase contrast microscopy movies depicting examples of cell twisting during division in multiple *ftsW**I\** Δ*fzlA* cells. As indicated, 5 minutes elapse between frames. Video playback is 10 frames per second. Strain key (*Caulobacter crescentus*): *ftsW**I\** Δ*fzlA* (EG2170).

## Supplementary Tables

**Supplementary Table 1:**

Tn-Seq data and analysis for WT vs. *ftsW**I* Caulobacter* genes. Columns WT 1, WT 2, WI 1, and WI 2 contain the number of unique transposon insertions in each gene in each replicate of WT or *ftsW**I\** (WI) strain transposon insertion library. These values were used to determine the log_2_ fold-change (logFC), log counts per million reads (logCPM), PValue, false-discovery rate (FDR), and negative log_10_ of the FDR for each gene in WT versus *ftsW**I\**. Genes are ordered by significance (neglog(FDR)).

**Supplementary Table 2:**

List of plasmids used in this study.

**Supplementary Table 3:**

List of strains used in this study.

